# Variability of functional and biodiversity responses to perturbations is predictable and informative

**DOI:** 10.1101/2022.06.20.496833

**Authors:** James A. Orr, Jeremy J. Piggott, Andrew L. Jackson, Michelle C. Jackson, Jean-François Arnoldi

**Affiliations:** Department of Zoology, University of Oxford, Oxford, UK; Zoology, School of Natural Sciences, Trinity College Dublin, Dublin, Ireland; Centre National de la Recherche Scientifique, Experimental and Theoretical Ecology Station, Moulis, France

**Keywords:** Ecology, Community Metrics, Aggregate Properties, Ecosystem Function, Diversity, Richness, Response Diversity, Multi-Functionality

## Abstract

Perturbations such as climate change, invasive species and pollution, impact the functioning and diversity of ecosystems. But because there is no unique way to measure functioning and diversity, this leads to a ubiquitous and overwhelming variability in community-level responses, that is often seen as a barrier to prediction in ecology. Here, we show that this variability can instead provide insights into hidden features of a community’s functions and responses to perturbations. By first analysing a dataset of global change experiments in microbial soil systems we show that variability of functional and diversity responses to a given perturbation is not random: aggregate properties that are thought to be mechanistically similar tend to respond similarly. We then formalise this intuitive observation to demonstrate that the variability of community-level responses to perturbations is not only predictable, but that it can also be used to access hidden and useful information about population-level responses to perturbations (i.e., response diversity and scaling by species biomass). Our theory offers a baseline expectation for the variability of community-level responses to perturbations and helps to explain the complexity of ecological responses to global change.

**Significance Statement:** Measures of biodiversity and ecosystem functioning show highly variable responses to a given perturbation. This variability is traditionally thought of as reflecting our inability to predict ecological responses to global change. Our work, however, finds that variability of community-level responses is itself predictable and can even be used to gain insights about how species respond to perturbations and collectively contribute to ecosystem functions.

## 1 Introduction

Understanding the behaviour of community-level features of ecosystems is a major goal of ecology. The importance and origin of species diversity was a central theme of late 20^*th*^ century ecology (Elton, 1958; Hutchinson, 1959; MacArthur, 1965; May, 1972), which led to a proliferation of metrics to define and measure diversity based on the richness, evenness and rarity of species (Simpson, 1949; Shannon & Weaver, 1963; Chao, 1987; Hill, 1973; Hurlbert, 1971). Since then, understanding how species collectively perform a function has become a prominent area of research (Ehrlich & Wilson, 1991; Tilman, Wedin, & Knops, 1996; Yachi & Loreau, 1999; Srivastava & Vellend, 2005), with clear implications for our understanding of concrete issues regarding productivity, carbon sequestration, pollination, or nutrient cycling of natural or engineered ecosystems. In light of rapid anthropogenic global change, there is currently increased focus on understanding how aggregate ecological properties will respond to perturbations such as land-use change, invasive species, climate change and pollution (Dornelas et al., 2014; Supp & Ernest, 2014; Paine, Tegner, & Johnson, 1998; Zhou, Wang, & Luo, 2020).

Ecologists are very aware that different aggregate properties, such as diversity metrics or ecosystem functions, describe very different aspects of communities and may thus respond in completely different ways to a given environmental perturbation. For instance, the many different diversity metrics employed by ecologists describe different statistical properties of the structure of communities, and have different purposes (Roswell, Dushoff, & Winfree, 2021; Magurran & McGill, 2010). If a perturbation caused the extinction of the rarest species while making the overall abundance more evenly distributed across surviving species, species richness would decrease, but a diversity metric more sensitive to common species (e.g., Simpson’s index) would potentially increase. Similarly, ecosystem functioning can be measured in a myriad of ways. Some functions, such as biomass production or respiration, are *broad* functions: they are performed by most or all species in a community. Other functions, such as the breakdown of specific chemicals or the production of specific enzymes, are *narrow* in the sense that they require the presence of particular species, or combinations of species, to be performed (Rivett & Bell, 2018; Hector & Bagchi, 2007). The great variety of ecosystem functions – in what they do, how broad or narrow they are, how species contribute to them, and how they respond to perturbations – has motivated the rapid development of multifunctional ecology where multiple functions are considered at once to more accurately characterize the state of an ecosystem (Manning et al., 2018; Delgado-Baquerizo et al., 2017; Giling et al., 2019). That being said, ecological systems are already complex entities, and on the surface, having to focus on this extra variability in how different functions and diversity metrics respond to perturbations, may seem like a dead-end, unlikely to help improve synthesis and prediction.

Here we propose that this is not the case, that on the contrary, much can be learned from the variability of functional and biodiversity responses to perturbations. To make this point we first analysed a dataset of global change experiments conducted in microbial soil systems (Box 1, Fig. 1). Focusing on three diversity metrics, two broad ecosystem functions, and eight narrow ecosystem functions, we analysed patterns of mismatches between observations. Concretely, we looked at the proportion of cases where one aggregate property responds negatively to a perturbation while the other responds positively to it. As expected, there was a great degree of variability in responses to perturbations. We found, however, that this variability is not random, but instead shows a recognizable degree of structure. Aggregate properties that are thought to describe ecosystems in similar ways (e.g., production of beta-xylosidase and production of cellobiohydrolase, enzymes that contribute to carbon cycling) have a lower proportion of mismatches than would be expected by chance (modules of blue squares, Fig. 1A). On the other hand, diversity metrics and ecosystem functions seem to systematically differ in how they respond to perturbations (dominance of red squares between diversity and ecosystem functions, Fig. 1A). Our intuitions about how mechanistically similar community aggregate properties are (i.e. how we ordered the observations Fig. 1A) thus seems to be a useful starting point for understanding the variability of their responses to perturbations, as it revealed clear patterns – some easy to interpret, and some more intriguing (such as the fact that functions and diversity tend to respond in opposite ways).

**Figure 1:**
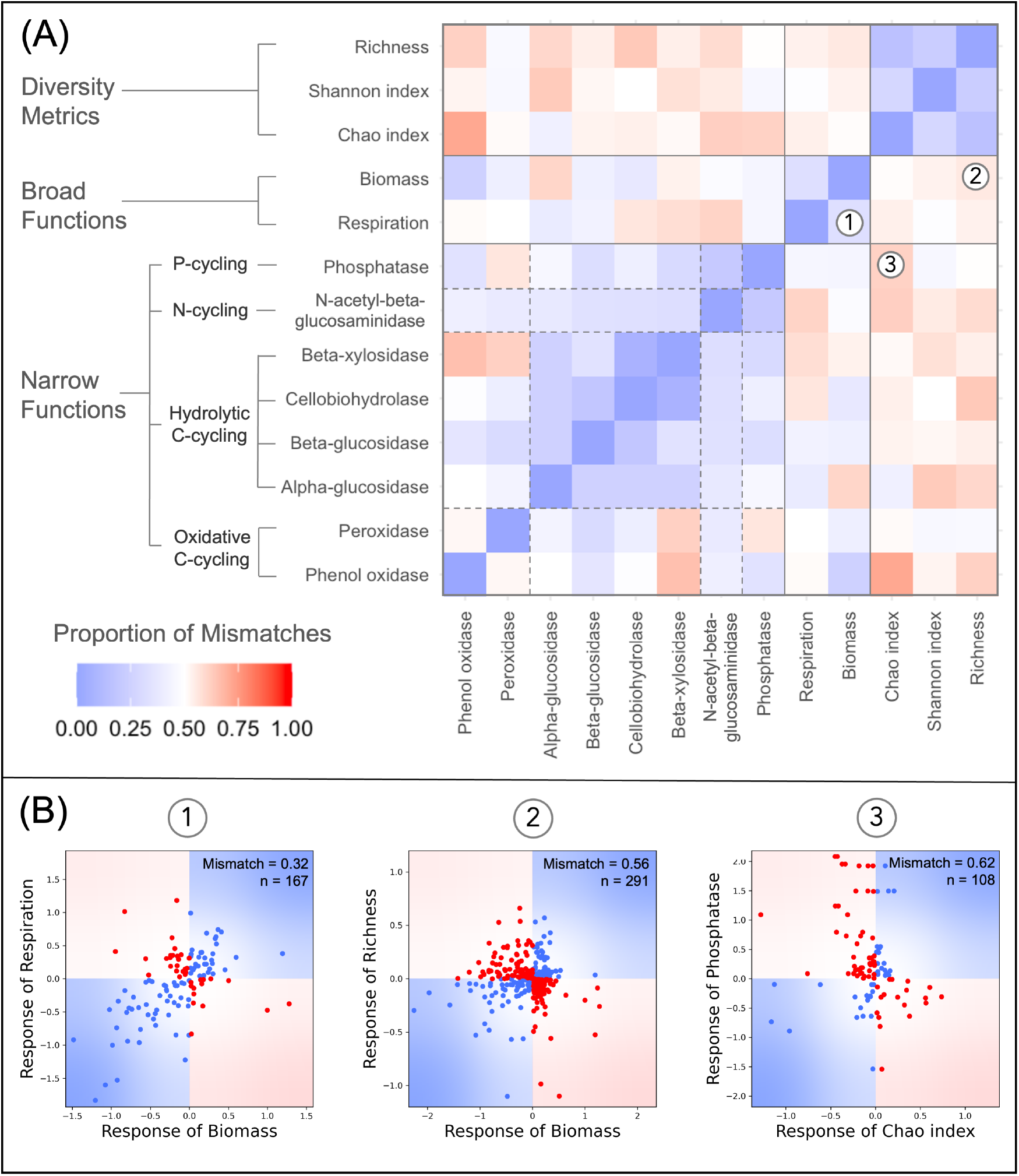
Synthesis of global change experiments in microbial soil systems quantifying variability of functional and biodiversity responses to perturbations. **(A)** The proportion of qualitative mismatches (one responds positively and the other negatively) between thirteen aggregate property including three measures of diversity (richness, Shannon index, Chao index), two broad ecosystem functions (biomass and respiration), and eight narrow ecosystem functions subdivided into P-cycling enzymes (phosphatase), N-cycling enzymes (N-acetyl-beta-glucosaminidase), hydrolytic C-cycling enzymes (beta-xylosidase, cellobiodydrolase, beta-glucosidase, alpha-glucosidase), and oxidative C-cycling enzymes (peroxidase, phenol oxidase). **(B)** Correlations between the response ratios of three specific pairs of aggregate properties: (1) biomass and respiration, (2) biomass and richness, (3) Chao index and Phosphatase. Points that fall in the blue areas of the plots are cases when the two metrics responded in the same way to a perturbation in a given experiment, while points that fall in the red areas are cases when there were qualitative mismatches between observations. The proportion of points that are red in these figures corresponds to the proportion of mismatches reported in (A).

Motivated by the findings of this empirical synthesis, we propose a theory that helps us understand, predict, and gain useful information from the variability of functional and diversity responses to perturbations. To do so, we convert the ecological problem into a simpler geometric one by representing perturbations as displacement vectors and community aggregate properties as directions in community state space. The central ingredient of our theory turns out to be a precise definition of collinearity between two aggregate properties which quantifies their similarity and predicts whether they will respond to a perturbation in the same way. This prediction assumes a high response diversity at the species level, and depends on how species’ responses scale with their biomass. Here, coarse-grained assumptions about population-level responses is used to better understand functions. Conversely, we show that with some knowledge of the aggregate properties used to observe a perturbation, the variability of their responses can be leverage to gain information about response diversity and biomass scaling at the population level. Our theory takes an accepted but seemingly inconvenient truth, that there is substantial variability between community-level observations, and uses it to gain insights about how ecosystems function and respond to perturbations.

**Box 1: Synthesis of empirical data**

To quantify the variability of functional and biodiversity responses to perturbations we analysed a dataset of global change experiments conducted in microbial soil systems (Zhou et al., 2020). This dataset contained 1235 perturbations from 341 publications. Perturbations included warming, elevated carbon dioxide levels, altered precipitation, nutrient enrichment, land-use change, or combinations of these factors. The effect of each perturbation in a given experiment was quantified using the natural logarithm-transformed response ratio:

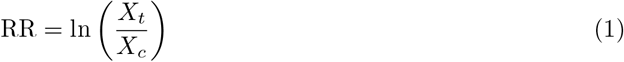

where *X*_*t*_ and *X*_*c*_ are the means of the treatment and control groups for a given aggregate property. Each individual perturbation was quantified using multiple aggregate properties covering a wide range of ecosystem functions and measures of diversity.

We focused on aggregate properties where all pairs had at least 10 observations in the dataset so that the proportion of mismatches between them could be estimated with some robustness. This gave us thirteen aggregate properties including three measures of diversity (richness, Shannon index, Chao index), two broad ecosystem functions (biomass and respiration), and eight narrow ecosystem functions subdivided into P-cycling enzymes (phosphatase), N-cycling enzymes (N-acetyl-beta-glucosaminidase), hydrolytic C-cycling enzymes (beta-xylosidase, cellobiodydrolase, beta-glucosidase, alpha-glucosidase), and oxidative C-cycling enzymes (peroxidase, phenol oxidase).

This list of aggregate properties was sorted *a priori* based on intuitions about their underlying mechanisms (grouped by diversity metrics, broad functions and narrow functions based on Zhou et al. (2020)) and a heatmap was made to visualize the proportion of mismatches between each pair (Fig. 1A). If the variability between aggregate properties was just random (i.e., if the heatmap was all white or just showed random distributions of red and blue) there might not be much more to say, but if the heatmap showed some structure there could be useful information to gain from the variability. Indeed, the modularity of the heatmap shows that aggregate properties that are thought to be similar tend to respond to perturbations similarly (e.g., relatively low proportion of mismatches – ranging from 0.16 to 0.28 – between measures of diversity). Conversely, groups of aggregate properties that describe different aspects of a community can systematically differ in their responses to perturbations (e.g., abundance of red between diversity metrics and ecosystem functions, with proportion of mismatches going as high as 0.73).

We will return to these empirical results in the discussion after we have outlined our geometric approach for quantifying the notion of similarity between aggregate properties. In fact, we can use our theory to reinterpret these empirical data to gain useful insights into how the perturbations in these experiments impacted these communities and also into how these communities collectively contribute to the different ecosystem functions. All data and code used in this synthesis are available at https://github.com/jamesaorr/community-properties.

## 2 Methods

### 2.1 Geometric Approach

To understand what can be learned from the variability of aggregate properties’ responses to perturbations, we transpose the ecological problem to a more abstract, but simpler, geometric setting (described more formally in Box 2).

First, we consider the effects of perturbations on populations as displacement vectors in the ecosystem’s state-space, where axes report the biomass of all constituent species (Fig. 2A). This vector is the difference between initial and perturbed states. It encodes the response to the perturbation at the population level. We then see ecosystem functions as positive directions in this same state space (Fig. 2B). Total biomass for example is the sum of all the species’ biomass and its direction lies exactly between all the axes, giving equal weight to all species. Other functions may not be influenced by the biomass of all species equally. In the hypothetical example shown in Fig. 2B, general decomposition is slightly more sensitive to the biomass of fungi than to the biomass of bacteria, plastic decomposition is primarily carried out by bacteria, and chemical production is primarily carried out by fungi. In general, a positive direction is spanned by a vector of positive values representing the per-capita contribution of each species to the function of interest. The “broadest” function, total biomass, is made up entirely of ones. The “narrowest” functions, are made up entirely of zeroes, except on the entry associated to the only contributing species (Rivett & Bell, 2018).

**Figure 2:**
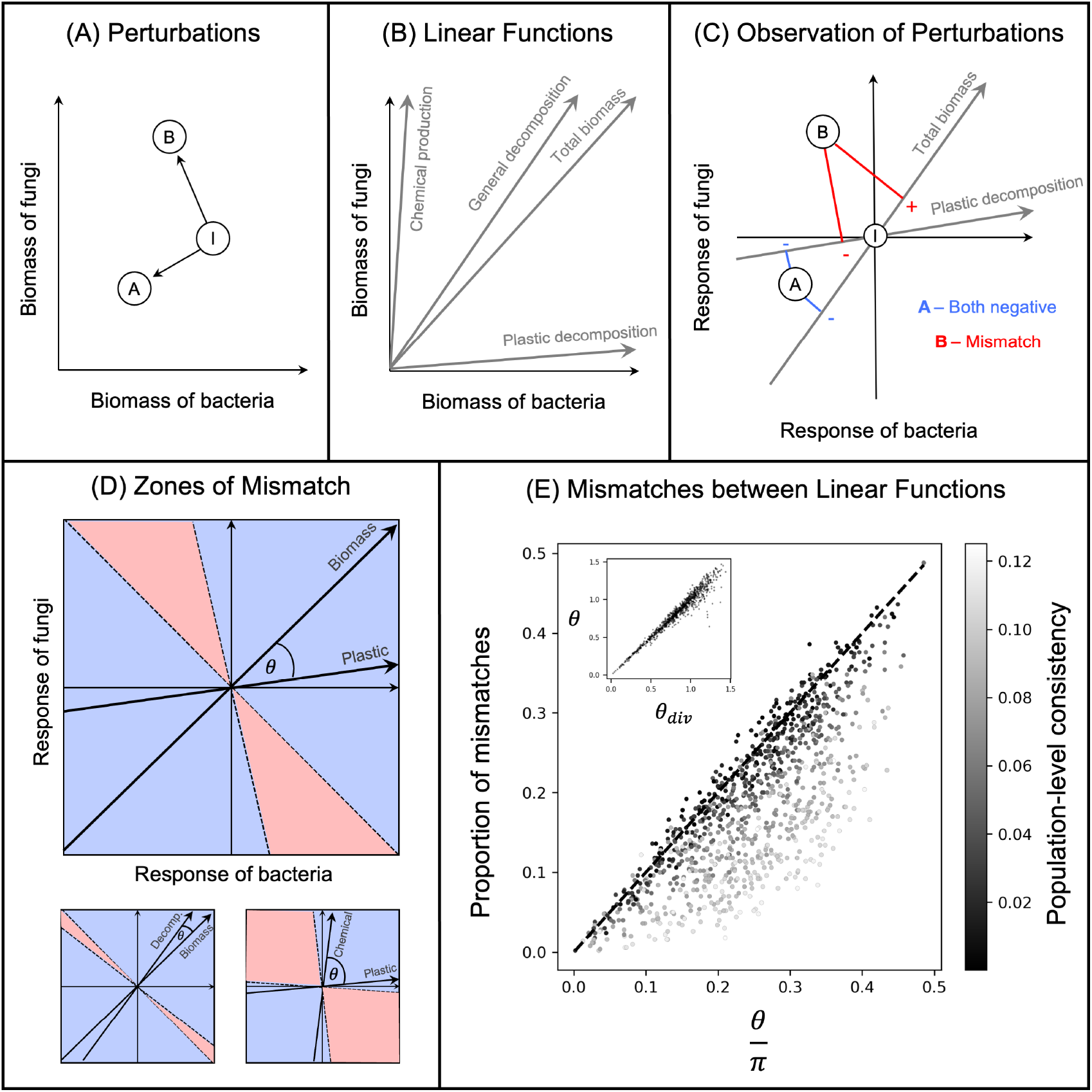
**(A)** Perturbations can be viewed as displacement vectors in community state-space. Here a community of bacteria and fungi is impacted by two perturbations, represented by the black arrows from the initial state of the community (*I*) to the points *A* and *B*. **(B)** Measures of ecosystem function can be represented as positive directions in this state-space. **(C)** Perturbed states are plotted in a space where the initial state of the community is at origin and each axis describes the response of each species to a perturbation. Here the displacement vectors associated with the perturbations *A* and *B* from (A) are projected onto the linear functions representing total biomass and plastic decomposition. For *A*, both functions observe negative responses. However, for *B* there is a mismatch in the observations of the functions: total biomass responds positively while plastic decomposition responds negatively. **(D)** For two functions, the zones of mismatches in their observations can be found by drawing lines perpendicular to the functions that go through the origin. Aggregate properties will observe different responses for perturbations that fall between these lines (i.e., in the red zones). The angle between the two functions determines the size of the zones of mismatches. **(E)** Over many *in silico* perturbation experiments, the proportion of mismatches between linear functions can be predicted by the angle between them in radians (*θ*) divided by the number *π*. When perturbations are consistent at the population level there are less mismatches than predicted as perturbations tend to fall in the mostly positive or mostly negative areas of state-space, which happen to overlap with the zones of consistent observations for linear functions (i.e., blue zones in (D)). The inset shows how the angle between two functions can be predicted using their diversities (Eq. (3)).

Next, we combine these two levels of abstraction to model how functions “observe” perturbations. We recenter the state space so that the axes now represent the response of each species, with the origin consequently being the initial state of the community (Fig. 2C). Projecting the displacement vector (multi-dimensional) onto the direction of an ecosystem function (one dimension) gives the “observation” of that function. For each function, drawing a line through the origin and perpendicular to the direction of the function delineates two zones. One where the projection is negative, and thus the function observes a negative response and the other where the projection is positive and thus the function observes a positive response. If the two directions associated to the two functions are not perfectly collinear, there will be zones of state-space where responses to perturbations will be qualitatively different when observed by one function or the other. These zones are the two symmetrical cones centered on the origin, formed by the delineation lines of the functions, perpendicular to their respective directions (red zones in Fig. 2D). The larger the angle between two functions, the larger the zones of mismatches. Consequently, if species’ responses were random and unbiased, the probability of finding a qualitative mismatch between two functions is:

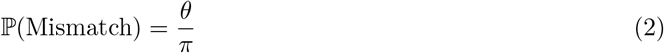

where *θ* is the angle between the two functions measured in radians. This collinearity of functions allows us to quantify their similarity. As the exact species contributions to a function may be challenging to acquire in empirical data, it is noteworthy that the angle between two functions can be approximated using only the knowledge, or estimation, of their respective broadness (Eq. (13) in Box 2). Indeed, in a community of *S* species and functions *f* and *g*:

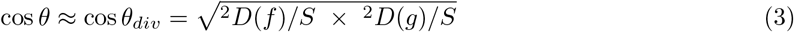

where 1*/S* ≤ ^2^*D*(*f*)*/S* ≤ 1 is the broadness of the function *f* (same for function *g*), defined here as the Gini-Simpson diversity index (Hill, 1973) of the vector of species contributions to the function, and normalized by species richness *S*. Expression (3) quantifies the intuitive expectation that two broad functions ought to be highly collinear, whereas two narrow functions can be independent (i.e. orthogonal to one another) if they are not performed by the same set of species.

Our final level of abstraction is the realization that measures of diversity, which are highly non-linear functions of species biomass (in the mathematical sense of a function of variables, not in the sense of ecological functioning), can still be placed into this geometric setting by considering their (state-dependent) gradients. The gradient of a diversity metric is a state-dependent vector encoding its sensitivities to small variations in each species biomass. Importantly, gradients of diversity metrics span non-positive directions in state-space because increasing the biomass of some species (the more abundant ones) decreases diversity.

### 2.2 Simulation toy model for perturbation experiments

To test, explore and illustrate the geometrical ideas outlined above, we conducted basic numerical experiments where ecological communities were perturbed and their responses were observed using different aggregate properties. We did not ask of our simulations to have complex, realistic underpinnings. We simply defined a protocol to generate a wide range of initial and perturbed states, and a wide range of aggregate properties (representing ecosystem functions or diversity measures) that we then used to quantify the ecosystem-level impacts of the perturbations.

Initial states were vectors of length *S* (chosen uniformly between *S* = 20 and *S* = 100) whose elements *N*_*i*_ are the initial species abundance or biomass. Those were drawn from log-normal distributions with zero mean and standard deviation (uniformly chosen between 1/2 and 2), thus generating a wide range of communities while also mimicking empirical abundance distribution patterns. For each initial state, 500 perturbations were generated as vectors Δ*N* of length *S* (perturbed states are *N* + Δ*N*) whose elements Δ*N*_*i*_ were generated in the following way. First, for each species we drew a value *x*_*i*_ from a normal distribution with unit standard deviation and mean *μ*. For a given initial state, *μ* is a fixed value uniformly chosen between -0.3 and 0.3. It determines the qualitative consistency of population-level responses (more on this below). We then normalized the set of values 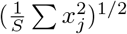, which gave us a set of values *y*_*i*_ that we used to define the actual response of species as

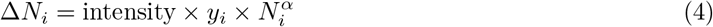

For a given perturbation, its intensity was drawn uniformly between 0 and 0.1. We also allowed the impacts of perturbations to scale with the initial abundance (or biomass, in this toy model there is no difference) of species. For each perturbation, the scaling exponent (*α*) was either 0 (no scaling, Figs. 2-3) or uniformly chosen between 0 and 1 (Figs 4-5). When *α* = 1, the population-response to the perturbation is, on average over the community, proportional to the species initial biomass. The other basic population level-feature that we considered is the response consistency (i.e., whether the perturbation impacted most species positively or negatively). As mentioned above, this feature is set by the parameter *μ*. Indeed, if we define the population-level response consistency as

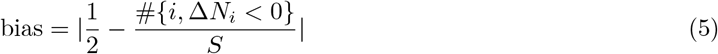

(# denotes the number of elements in a discrete set, here the set of species whose abundances are reduced by the perturbation); then, the expected fraction of negative population responses in the above expression is simply Φ(−*μ*) where Φ(*x*) the cumulative function of a standard normal distribution.

Ecosystem functions, which we used to “observe” the ecosystem-level response to perturbations, were represented by positive directions in an *S*-dimensional space, spanned by vectors ***φ*** whose elements *φ*_*i*_ represent species’ per-capita functional contributions. For a given state *N* = (*N*_*i*_), its level of functioning is then *f* (*N*) = *φ*_*i*_*N*_*i*_ (see Box 2). The per-capita contributions *φ*_*i*_ were drawn from a log normal distribution with a standard deviation uniformly chosen between 0 and 1.3. When the standard deviation was small, the functions were broad as the per-capita contributions of each species were similar. When the standard deviation was large, however, the functions were more narrow, with a large variation in the per-capita contributions of each species to the function.

Diversity metrics were taken from the family of Hill diversity that define the effective number of species as:

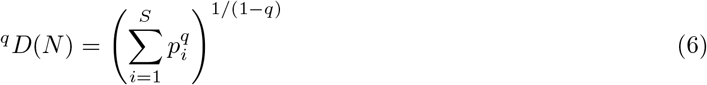

where *S* is richness, *p*_*i*_ is the relative abundance (or biomass) of species *i* and *q* is the hill number that determines the sensitivity of the diversity index to rare or to abundant species. This general equation encompasses species richness (*q* = 0), the Shannon index (*q* = 1) and the Gini-Simpson index (*q* = 2) (Roswell et al., 2021; Hill, 1973; Shannon & Weaver, 1963; Simpson, 1949). In simulations we chose the indices *q* between 0 and 5, thus spanning a wide range of diversity measures. To apply our geometrical theory to diversity observations we considered the directions spanned by their gradients (the vector of partial derivatives 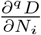), evaluated at the initial state, which take the form ^*q*^***φ*** = (^*q*^*φ*) with

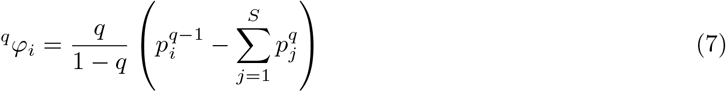

For each perturbation experiment, and each pair of aggregate properties, *f, g* (either two positive linear functions, two diversity measures, or one of each), we checked the consistency of their responses. That is, we looked at the sign of *f* (*N* + Δ*N*) − *f* (*N*) and compared it to the sign of *g*(*N* + Δ*N*) − *g*(*N*). If they do not coincide, there is a qualitative mismatch between the two ways of observing the ecosystem’s response to the perturbation.

For Fig. 2E, 1000 communities were generated and each one experienced 500 different perturbations. For each community, two ecosystem functions of varying broadness were generated and used to observe the community-level responses to the perturbations. The angle between the directions defined by the functions was calculated, divided by *π* (Eq. 2), and plotted against the realised proportion of mismatches over the 500 perturbations, while recording the relative deviation from the prediction. The angle between each pair of functions was also estimated using only the knowledge of their broadness based on Eq. (3). For Fig. 3, two diversity indices with different *q* numbers were generated to observe the responses of a given community to 500 perturbations. The angle between their gradients divided by *π* was used to predict the proportion of mismatches. For Fig. 4A, total biomass (positive direction whose elements are all 1) and Hill-Simpson (^2^*D*) were used to observe the ecosystem-level responses to the perturbations. The effective angle between total biomass and the state-dependent gradient of the diversity index, based on Eq. 15, was calculated, divided by *π*, and plotted against the actual proportion of mismatches. For Fig. 4B, the actual responses of total biomass and diversity (*q* drawn from a random uniform distribution between 0 and 2) were plotted for 1000 perturbations that were scaled by biomass to a certain degree (*α* drawn from a random uniform distribution between 0.5 and 1). All data and code used for the simulations are available at https://github.com/jamesaorr/community-properties.

**Figure 3:**
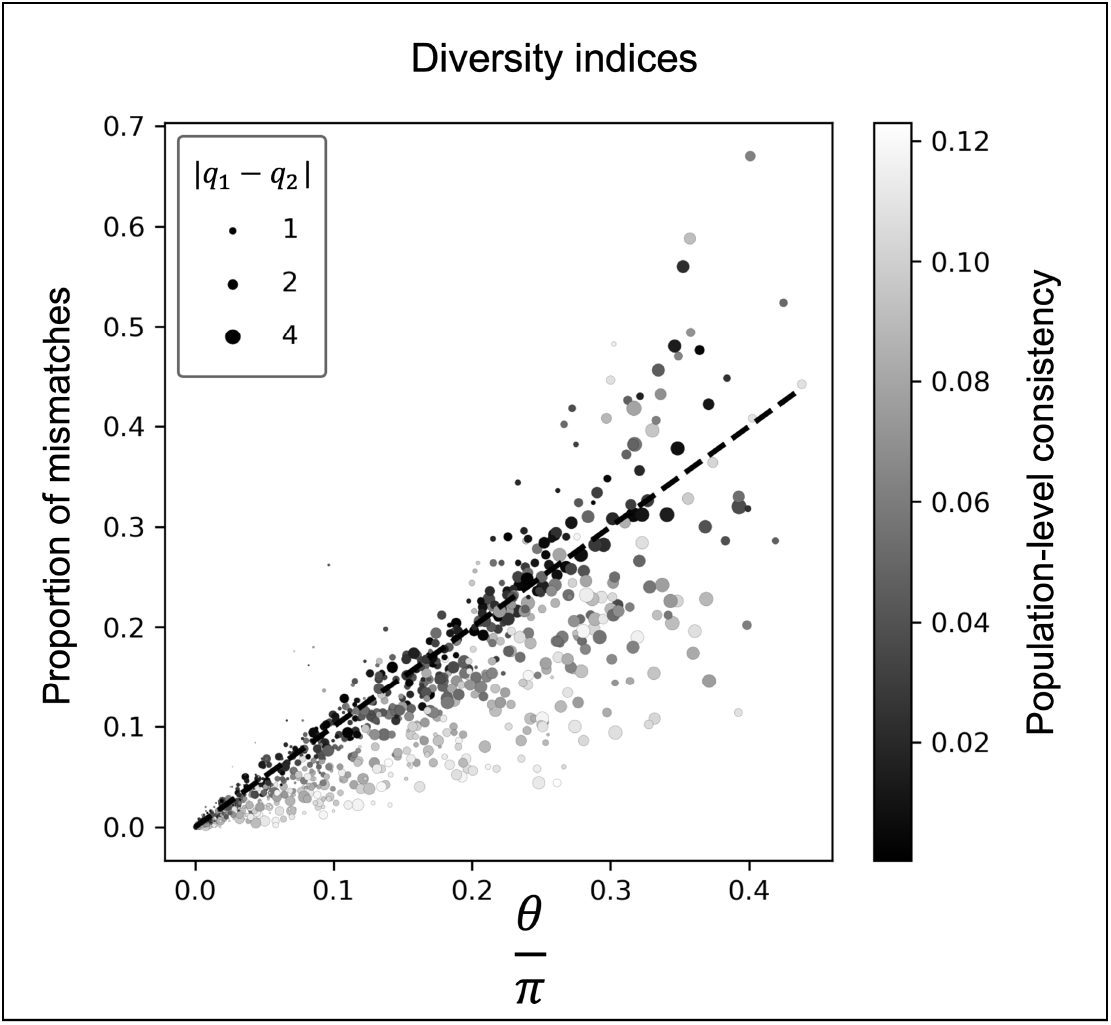
The proportion of mismatches between two diversity metrics is predicted by the angle between their gradient vectors. All diversity metrics came from the family of Hill diversity indices so increasing the difference between the *q* values of diversity indices (size of the points) increases the angle between their gradients, which therefore increases the proportion of mismatches.

**Box 2: Formalizing the variability of observed responses to a perturbation**

We formalize the process of observing the ecosystem level impact of a given perturbation, based on aggregate features of functioning or diversity. Our goal is to explain what controls the probability that two scalar observations of the same perturbed ecosystem give opposite results. Here bold symbols denote *S*−dimensional vectors, where *S* is the richness of the community. Let ***N*** ^*c*^ be the initial state of a community: the vector of species biomass prior the perturbation. Let ***N*** ^*p*^ be the perturbed community state. The observed response, quantified via an ecosystem function *f* (***N***), is

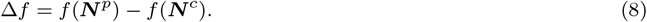

For a linear function there exists a constant *f*_0_ (because we will consider changes in functioning, and not absolute levels of functioning, this constant will play no role in what follows) and a vector ***φ*** –the gradient– such that

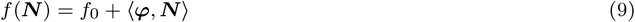

with ⟨·, ·⟩ the scalar product of vectors. The elements of the gradient vector ***φ*** encode the per capita contribution of species to the function. For us it will not matter what those exact contributions are, only relative species contributions which determine the *direction* spanned by the vector ***φ***. A positive function is such that the elements of the gradient are positive. If we rewrite the response of the function, we get that

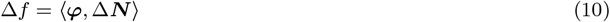

where Δ***N*** = ***N*** _*p*_ − ***N*** _*i*_ is the vector of population-level responses. For non-linear functions the (state dependent) gradient vector can be computed as 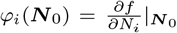. In this case, expression (10) will be an approximation, accurate for weak perturbations. Now, for two functions, *f, g* associated to two directions spanned by the two gradient vectors ***φ*** and ***ϕ***, we define their collinearity as the angle 0 ≤ *θ <* 2*π* whose cosine is

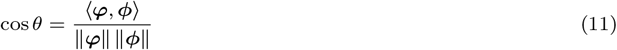

where ∥·∥ denotes the Euclidian norm of vectors. A graphical argument (Fig. 2D) tells us that the fraction of perturbation vectors Δ***N*** that will lead to a mismatch between the observations of *f* and *g* is

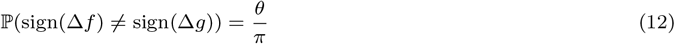

In such cases, one of the functions will observe a positive response, while the other function will observe a negative response. For random positive directions, we can evaluate the cosine of the angle based on a notion of functional broadness. Indeed, given a random choice of positive functions

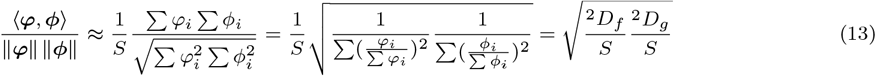

Where ^*q*^*D* denotes Hill’s diversity index. We will call the fraction 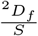 the *broadness* of the function *f*, which is maximal (and equal to one) if all species contribute equally to the function (i.e. total biomass).

We can modify the above theory to account for an additional piece of population-level information in the form of a biomass scaling of population-level responses. It is indeed reasonable that more abundant species will, in absolute terms, show a larger response to perturbations. For some scaling exponent *α* ≥ 0, if we denote Λ the diagonal matrix whose elements are the species biomass prior to the perturbation, we may assume that the perturbation displacement vector takes the form Δ***N*** = Λ^*α*^**Δ**. We then have that

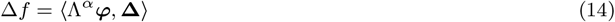

the relevant angle to consider then becomes

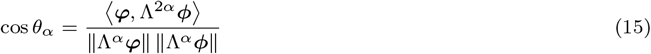

giving the fraction of rescaled vectors **Δ** that would lead to a qualitative mismatch.

## 3 Results

### 3.1 Mismatches between functions

As expected from our geometrical arguments, over many simulated perturbation experiments the proportion of mismatches between the observations of two ecosystem functions is predicted by their collinearity, as defined by the angle between their respective directions (Eq. 2, Fig. 2E). We also confirm that this angle can be estimated using only the functions’ respective broadness (Eq. 3, Fig. 2E inset). However, we see that the probability of mismatches is accurately predicted only when perturbations are unbiased at the population level; that is when approximately half the species show positive responses and half the species show negative responses. If population-level responses are biased towards positive or negative, our prediction overestimates the proportion of mismatches (Fig. 2E, Fig. 5A). This effect occurs because when perturbation vectors are mostly positive or negative, they tend to fall in the areas where positive linear functions must always observe the same responses (top right and bottom left quadrants in Fig. 2D). This could be seen as a limitation, but it actually provides a useful link between mismatches in observations at the community level and information on population-level response diversity (Fig. 5A). Indeed, deviations from our predictions reveal a a degree of population-level consistency in response to the perturbation considered. Furthermore, the deviation from the expectation is enhanced when considering species rich systems (shown by the colour gradient in Fig. 5A). This effect of dimensionality can be understood semi-intuitively: a bias at the population level increases the probability of the community response to fall in the positive or negative quadrants (where all species respond in the same direction) in which case any positive function will necessarily respond in the same way. This probability is (1*/*2 + *b*)^*S*^ where *b* is the bias, and so we see that the effect of *b* is exponentially accentuated by species richness (Fig. 5A). Therefore, if we can make a prediction based on the knowledge of species contributions to functions, or at least their broadness (Eq. 3), deviations from our predictions together with knowledge of diversity can give us insights into the system’s response diversity, in the form of our measure of population-level consistency.

### 3.2 Mismatches between diversity metrics

Diversity metrics are non-linear functions, but from the angle between their gradients, we can still predict response mismatches. Here we compare metrics from the family of Hill diversity, which vary based on *q*, the hill exponent, which controls their sensitivity to common or rare species (Roswell et al., 2021). Consequently, the difference in the *q* values between two diversity metrics correlates to the size of the angle between their gradients (Fig. 3A). The consistency of population-level responses has a weaker effect here due to the fact that diversity metrics are associated to non-positive functions. Therefore, a bias towards positive or negative population-level responses does not necessarily translate as a bias at the level of diversity measures. In fact, the evenness of the biomass distribution of a community determines how similar gradients of diversity metrics are to positive functions. For uneven communities, increasing the biomass of most species will increase diversity so gradients will behave similarly to positive linear functions. However, for perfectly even communities, increasing the biomass of any single species will decrease diversity so gradients of diversity will effectively be negative functions of biomass.

### 3.3 Mismatches between function and diversity

The probability of mismatches between ecosystem functions and diversity metrics can also be predicted by considering the angle between the function and the gradient of the diversity metric (Fig. 3A)). As before, consistency of responses at the population level causes the prediction to overestimate the actual proportion of mismatches. However, we now see that the angle between these two aggregate properties can exceed 90 degrees leading to a systematic bias towards mismatches of observations. This intriguing result is connected to a second piece of population-level information: the scaling of perturbations by species biomass (Eq. (14) in Box 2).

When the effect of perturbations are larger for more abundant species (as *α* approaches 1), function and diversity are expected to observe qualitatively different responses (only the larger points are above the red line in Fig. 3C). If a perturbation causes the biomass of abundant species to decrease, total biomass will decrease but diversity will increase. If on the other hand, a perturbation causes the biomass of abundant species to increase, total biomass will increase but diversity will decrease. Indeed, how much a perturbation is scaled by biomass can be predicted based on the proportion of mismatches between function and diversity (Fig. 5B). The empirical data (Fig. 1B), can be replicated by simulating many perturbation experiments and comparing how function and diversity observe perturbations. To recreate the empirical results, where the proportion of mismatches between function and diversity exceeds 60%, perturbations must be scaled by biomass to a power greater than *α* = 0.5 (Fig. 3D).

## 4 Discussion

Variability of results, or “context-dependency”, is infamously pervasive in ecology (Catford, Wilson, Pyšek, Hulme, & Duncan, 2021). While this is partly what makes ecosystems so fascinating to study, it is also an obstacle to the synthesis of previous results and to the prediction of future impacts. Our research has focused on some of this variability – the variability between the responses of community aggregate properties to a given perturbation – and found that it is predictable and also a rich source information. Mismatches between the responses of different aggregate properties to a perturbation can give us previously difficult to access information about the aggregate properties themselves (e.g., similarity between ecosystem functions) and about how that perturbation is impacting the species that constitute the community (e.g., response diversity and biomass scaling). Ecological research is typically reductionist, using information about individuals and populations to understand communities and ecosystems (Loreau, 2010). Our work demonstrates the reverse approach by using information about communities to understand population-level responses. Before discussing the potential future directions this line of research could take, we should first return to the empirical synthesis presented in Box 1 to see what else can be said about these data in light of our theory.

### 4.1 Reinterpretation of Empirical Results

Upon first inspection of the empirical data, we could deduce from the modularity of the results (heatmap shown in Fig. 1 in Box 1) that aggregate properties that are known to be similar also tend to respond similarly to perturbations. This observation is certainly reassuring, as it confirms that mechanistic understandings at the chemical-level of microbial functions are consistent with community-level observations. But now that we have a quantitative way of measuring the similarity of aggregate properties (via their collinearity in state space) and we know about a connection to population-level behaviour (response diversity and biomass scaling), we can take our biological interpretations a few steps further.

Firstly, based on the systematic mismatches between diversity and function, we may deduce that the perturbations in these experiments must scale with species biomass, in the sense that species that initially represent a large proportion of the overall biomass also represent a large proportion of the variation in biomass caused by the perturbation. Indeed, we showed that diversity and function will respond to perturbations in qualitatively different ways only when the biomass scaling exponent (*α*) is above 0 (Figure 4). This tells us that the perturbations in these experiments have greater impacts on more abundant species. It is also tempting to say that the perturbations in these experiments have low levels of population-level consistency. We see that total biomass and respiration, which we might expect to be highly collinear as they are both broad functions, still show a relatively high proportion of mismatches (0.32). In fact, there are only few cases where aggregate properties show highly consistent responses to perturbations: only one of the 78 pairs of functions had less than 0.15 proportion of mismatches. Based on our geometrical arguments, if perturbations were even moderately consistent at the population level we would expect much lower levels of mismatches (Fig. 2). This interpretation, however, assumes that the functions are accurately represented by *positive* directions, meaning that all species contribute positively to the function. This may not be always be the case. Our theory can help to identify this.

**Figure 4:**
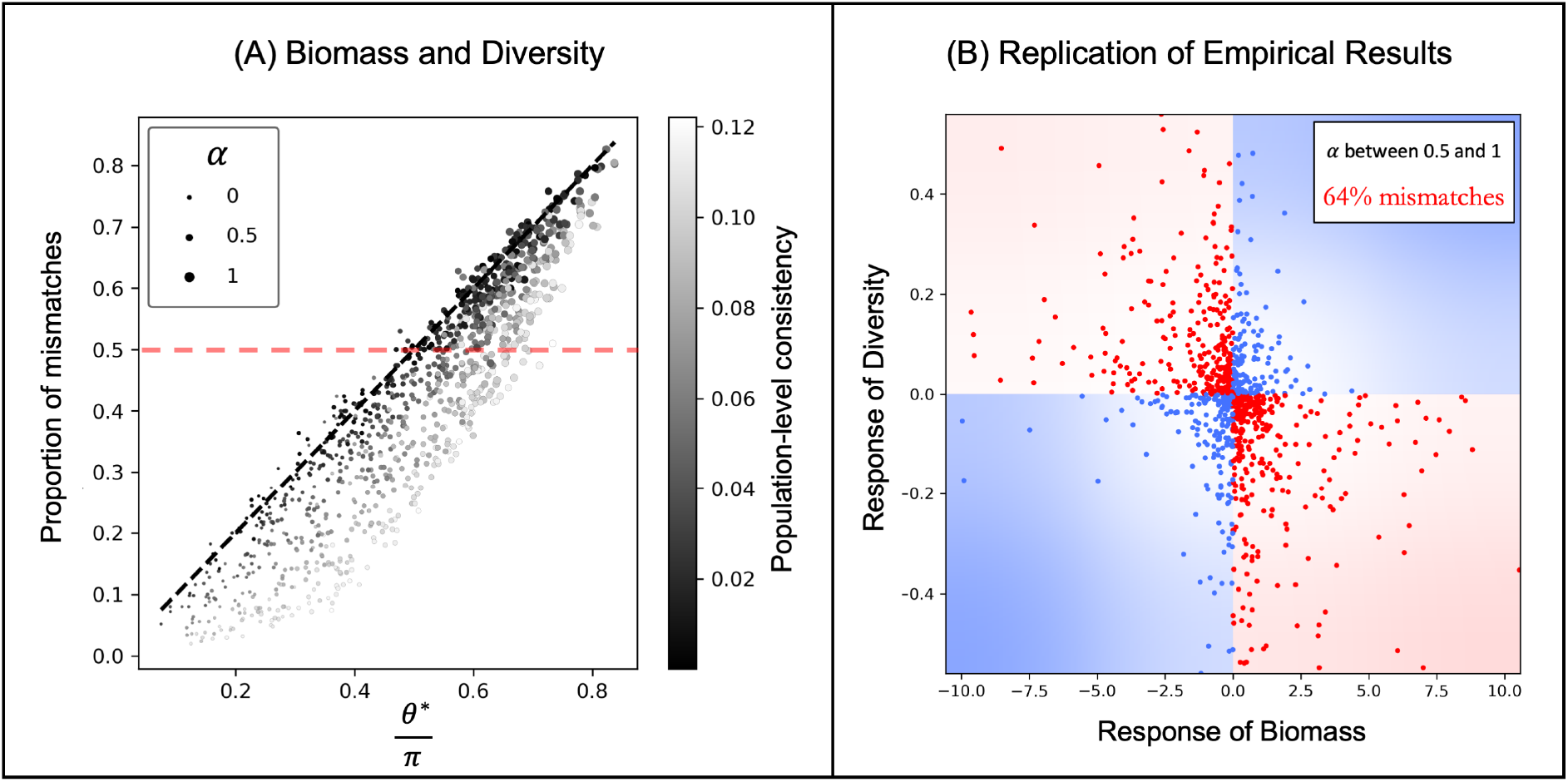
**(A)** Over simulated perturbation experiments, the angle between total biomass and the gradient of a diversity index (here *q* = 2) predicts the proportion of mismatches between them. Specifically, the relevant angle is between total biomass and the gradient of diversity after they have been scaled by the biomass of each species (*θ*^*^). When perturbations are scaled by species biomass (scaling exponent *α >* 0) total biomass and diversity can effectively become opposite functions. Points above the dashed red line show cases where there is a systematic mismatch in the observations of total biomass and diversity. **(B)** The empirical results from Fig. 1B can be recreated by simulating perturbation experiments where population-level responses to perturbations are relative to species’ biomass (specifically, *α* is drawn from a uniform distribution between 0.5 and 1).

**Figure 5:**
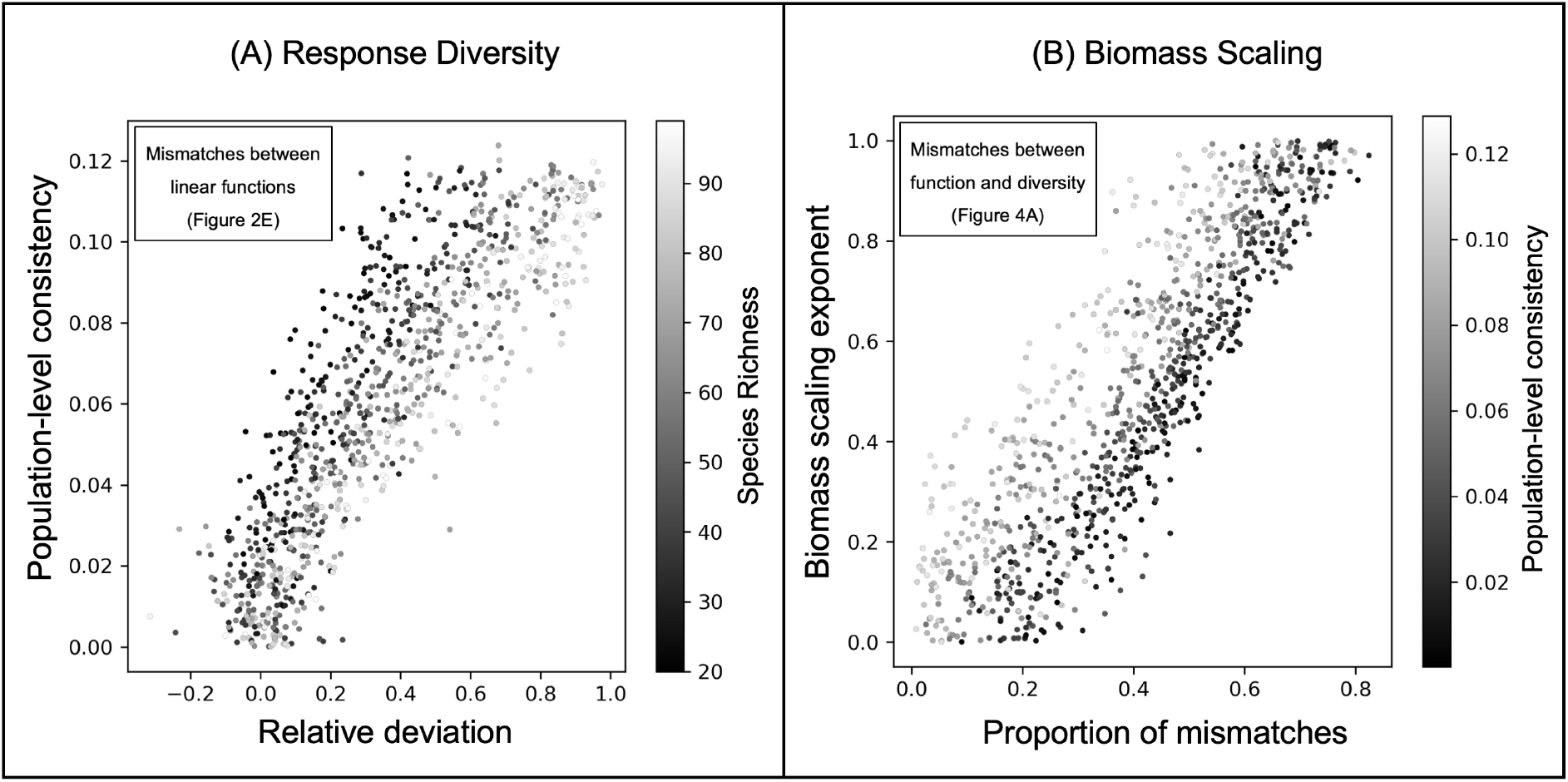
**(A)** A notion of the response diversity of a perturbation, the consistency of population-level responses, can be estimated from the relative deviation from our baseline expectation (Eq. 2). For a given amount of population-level consistency, increasing species richness will lead to a greater deviation from the expectation. Data is from the simulations presented in Fig. 2E. **(A)** How much a perturbation is scaled by biomass can be estimated from the proportion of mismatches between function and diversity. Data is from the simulations presented in Fig. 4A.

When functions can be represented by positive directions the proportion of mismatches cannot exceed 0.5 as the angle between their directions cannot exceed 90 degrees (see Fig. 2). In the empirical data, however, there are several pairs of functions (9/45) that show 0.5 proportion of mismatches, or more. This must mean that not all species are contributing positively to those functions. When some species, perhaps through their interactions with others, negatively contribute to an ecosystem function the angle between the directions associated to the functions can exceed 90 degrees. The directions then oppose each other which is how a systematic mismatches between the function’s responses can occur. For total biomass (which by definition is a positive function) to systematically differ to alpha-glucosidase (0.61 proportion of mismatches), it must be that some species in those microbial soil communities negatively contribute to the production of this enzyme. These insights demonstrate how our theory can be used to gain a richer understanding of the functioning of ecosystems and the perturbations that impact them.

### 4.2 Future Directions

Our geometric framework should be thought of as a null model for the variability between community-level responses to perturbations. It does not attempt to predict how specific aggregate properties will respond to specific perturbations. Instead, our theory can be used to test hypotheses. It can generate a non-trivial null expectation, based on the assumption that functions can be seen as directions (which amounts to assuming that per-capita contributions of species to functions are fixed). When this expectation is rejected by the data this tells us that something biologically interesting is going on. For example, we could imagine that when a pulse-type perturbation reduces total biomass of a community there would be a sudden increase in the production of enzymes related to rapid growth, which would result in systematic mismatches between measures of ecosystem function. Our theory offers a solid platform from which we can begin to explain these sorts of observations. More generally, it reveals two promising research avenues.

Our research illustrates that population-level responses to perturbations can be studied from the top-down by comparing the observations of different aggregate properties (Fig. 6A). What we called population-level consistency of a perturbation is closely linked to the concept of response diversity (Elmqvist et al., 2003). Response diversity – the variation between species responses to a perturbation – can be measured in different ways and is a key mechanism underlying ecological stability and the biological insurance hypothesis (Ross, Petchey, Sasaki, & Armitage, 2022; Loreau et al., 2021). Although the information we can gain using our geometric approach (i.e., the proportion of species responding positively or negatively – see Fig. 2) is a coarse measure of response diversity, it can be accessed by just comparing the observations of different aggregate properties rather than actually measuring each species’ response. Furthermore, whether the direct effect of a perturbation is proportional to the biomass of each species (which we quantify using the biomass scaling exponent, *α*) is another useful piece of information that can be gained using our top-down approach. This feature of perturbations controls the relative importance of rare or common species in determining the community’s temporal variability (“environmental perturbations” *sensu* Arnoldi et al., (2019)). Comparing multiple community-level observations (the more comparisons the better) allows us to describe these features of perturbations without ever having to collect information directly at the population level, which could therefore be an efficient and cost-effective tool for research synthesis and the analysis of biomonitoring data.

**Figure 6:**
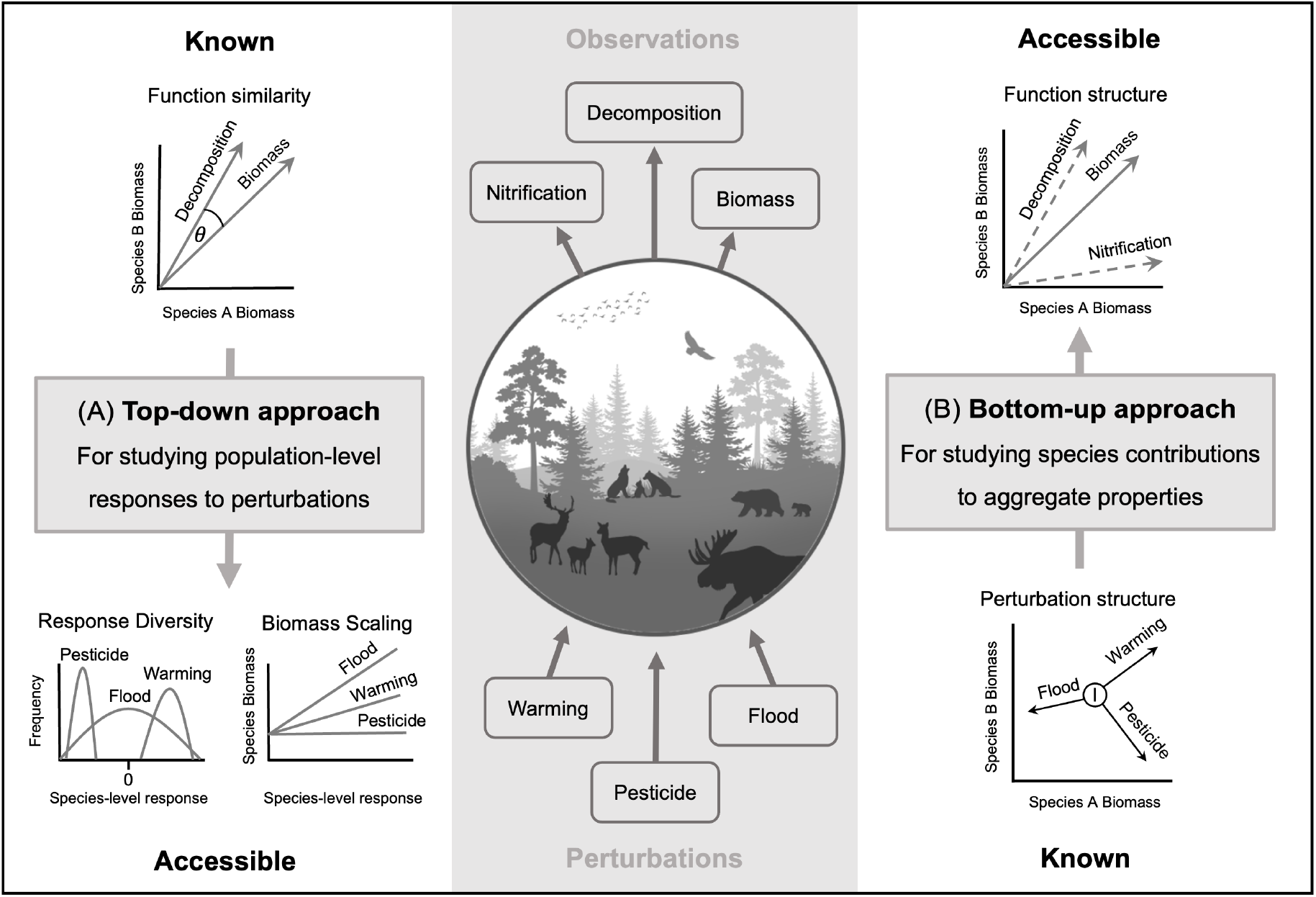
Overview of how our theory connecting variability of community-level responses with population-level information opens up two promising avenues of research. The three perturbations (warming, pesticide and flood) and the three aggregate properties (nitrification, decomposition and biomass) are used here for illustrative purposes only. **(A)** If the similarity of functions is known, features of perturbations (response diversity and biomass scaling) can be estimated using a top-down approach. **(B)** If a community is perturbed in known directions, species’ contributions to ecosystem functions can be estimated using a bottom-up approach.

But we can also flip this line of thought, which has been the dominant perspective of this article so far, to study species contributions to aggregate properties from the bottom up (Fig. 6B). If we wanted to know how a community of species performed some function, we could perturb the community in known directions and compare the unknown function’s responses to some known function (e.g. total biomass). Using perturbations as “probes” it would be possible to uncover the species’ unknown functional contributions (i.e., reconstruct the function’s direction in state space) by quantifying its similarity to the known function. Understanding the links between community composition and functioning has far reaching implications for many sectors including ecosystem management, agriculture, forestry and medicine (Diaz-Colunga, Skwara, Vila, Bajic, & Sánchez, 2022; Ross et al., 2021; Costea et al., 2018) and our approach contributes to the recent effort to study ecosystem functions in their natural context, in contrast to the traditional reductionist approach of using controlled experiments where populations or even organisms are studied in isolation (Bergelson, Kreitman, Petrov, Sanchez, & Tikhonov, 2021).

### 4.3 Conclusions

Given the generality of our theory, our results unsurprisingly touch many areas of contemporary ecology. For multifunctional ecologists, our work helps to explain how different functions can respond in different ways to global change (Giling et al., 2019). For ecologists interested in multiple perturbations, our theory can be used to understand variability in how community-level properties observe the interactions (antagonistic or synergistic) between perturbations (Orr et al., 2020). For biodiversity-ecosystem functioning research, the opposing responses of diversity and function to perturbations, which we show can often be expected, should be considered when understanding how perturbations influence biodiversity-ecosystem functioning relationships (Brose & Hillebrand, 2016). When studying complex systems such as ecosystems, it is important to have baseline expectations for their behaviour. We have found that the variability between community-level responses perturbations is not just a dead-end that limits synthesis and prediction in ecology. Instead, this variability is predictable and can be leveraged to gain useful information about species’ responses to perturbations and species’ contributions to ecosystem functioning. Our work provides a solid platform from which the complexity of community-level responses to anthropogenic global change can be better understood.

## Acknowledgements

JAO and MCJ were supported by a Natural Environment Research Council grant (NE/V001396/1). JAO was supported by an Irish Research Council Laureate Award (IRCLA/2017/112) and TCD Provost’s PhD Award held by JJP. JFA and ALJ were supported by an Irish Research Council Laureate Award (IRCLA/2017/186). JFA was supported by the TULIP Laboratory of Excellence (ANR-10-LABX-41).

